# Impact of genomic preselection on subsequent genetic evaluations with ssGBLUP - using real data from pigs

**DOI:** 10.1101/2021.06.18.449002

**Authors:** Ibrahim Jibrila, Jeremie Vandenplas, Jan ten Napel, Rob Bergsma, Roel F Veerkamp, Mario P.L Calus

## Abstract

**Background:** Empirically assessing the impact of preselection on subsequent genetic evaluations of preselected animals requires comparison of scenarios taking into account different approaches, including scenarios without preselection. However, preselection almost always takes place in animal breeding programs, so it is difficult to have a dataset without preselection. Hence most studies on preselection used simulated datasets, concluding that genomic estimated breeding values (GEBV) from subsequent single-step genomic best linear unbiased prediction (ssGBLUP) evaluations are unbiased. The aim of this study was to investigate the impact of genomic preselection (GPS) on accuracy and bias in subsequent ssGBLUP evaluations, using data from a commercial pig breeding program.

**Methods:** We used data on four pig production traits from one sire line and one dam line. The traits are average daily gain during performance testing, average daily gain throughout life, backfat thickness, and loin depth. As these traits had different weights in the breeding goals of the two lines, we analyzed the two lines separately. Per line, we had a reference GPS scenario which kept all available data, against which the next two scenarios were compared. We then implemented two other scenarios with additional layers of GPS by removing all animals without progeny either i) only in the validation generation, or ii) in all generations. We conducted subsequent ssGBLUP evaluations per GPS scenario, utilizing all the data remaining after implementing the GPS scenario. In computing accuracy and bias, we compared GEBV against progeny yield deviations of validation animals.

**Results:** Results for all traits in both lines showed marginal loss in accuracy due to the additional layers of GPS. Average accuracy across all GPS scenarios in both lines was 0.39, 0.47, 0.56, and 0.60 respectively for the four traits considered in this study. Bias was largely absent, and when present did not differ greatly among corresponding GPS scenarios.

**Conclusion:** As preselection generally has the same effect in animal breeding programs, we concluded that impact of preselection is generally minimal on accuracy and bias in subsequent ssGBLUP evaluations of selection candidates in pigs and in other animal breeding programs.

## Background

In animal breeding, parents of the next generation are often selected in multiple stages, and the initial stages of this selection are called preselection [1–3]. Selection candidates that survive preselection are called preselected animals [1–3], and those that do not are called preculled animals [3,4]. Preselection aims to reduce costs and efforts spent on animals that are not interesting for the breeding program, and achieves this by avoiding phenotyping or further testing of preculled animals. Due to introduction of genomic prediction [5], preselection is now mostly based on genomic estimated breeding values (GEBV) of young animals even before they have records of any traits. This type of preselection is called genomic preselection (GPS; e.g. [1,2]). The popularity of GPS is because genotyping is becoming cheaper by the day, and the reasonable reliabilities of GEBV (e.g. [6–8]). As genomically preculled animals have neither progeny nor records for some or all breeding goal traits, they are generally not included in subsequent genetic evaluations (i.e. genetic evaluations that come after preselection). GPS therefore decreases the amount of information available for subsequent genetic evaluations of preselected animals. Properly assessing the impact of preselection on subsequent genetic evaluation of preselected animals requires comparison of scenarios taking into account different approaches, including a scenario without preselection. Because in animal breeding programs preselection almost always takes place, it is difficult, if not impossible, to have a scenario without preselection. This is why most studies available on preselection used simulated datasets (e.g. [1,3,9,10]). Those studies have shown that when a subsequent genetic evaluation of preselected animals is done using pedigree-based best linear unbiased prediction (PBLUP), preselection results in accuracy loss and bias in the estimated breeding values (EBV) of preselected animals [1,3,9–12]. Some of these studies [9–12] further showed that the accuracy loss and bias caused by GPS can be avoided if the information on preculled animals that was utilized at preselection is included in subsequent PBLUP evaluations. On the other hand, our previous works [3,4] have shown that when the subsequent genetic evaluation is done with single-step genomic BLUP (ssGBLUP), genomic EBV (GEBV) of preselected animals are estimated without bias. We [4] further showed that to avoid GPS bias in subsequent ssGBLUP evaluation of preselected animals, genotypes of their preculled sibs are only needed if not all of their parents are genotyped.

In our previous works [3,4], being based on simulated datasets, preselection was the only possible source of bias in ssGBLUP evaluations. However, in real breeding programmes, other sources of bias in ssGBLUP evaluations may exist and are potentially difficult to control. Therefore, impact of preselection might be confounded by the impact of these other factors. These other possible sources of bias include, amongst others, inaccurate or incomplete pedigree [13], inaccurately estimated additive genetic (co)variances [13], and a reference population of selectively genotyped animals [14,15]. Although some ways to reduce the bias caused by these factors have been developed, the bias is usually not completely eliminated in evaluations using real data (e.g. [13–15]). This may explain the observation that in practice GEBV obtained from ssGBLUP evaluations are sometimes biased. The aim of this study was to investigate the impact of GPS on accuracy and bias in subsequent ssGBLUP evaluations, using data from a commercial pig breeding program in which preselection has taken place. To achieve this aim, we used the full dataset as control and retrospectively implemented additional layers of GPS. The additional layers of GPS were implemented by discarding animals that did not have progeny in the data. Since in the breeding program GEBV were used to select parents of next generations, discarding animals without progeny in the data can be considered as additional GPS. Then we compared results from subsequent ssGBLUP evaluations after these additional layers of GPS against results from ssGBLUP evaluation using the full available data. Our subsequent genetic evaluations only involved reevaluating preselected animals, either with or without preculled animals in the subsequent evaluations.

## Methods

### Data

We obtained pig production traits data on one sire line and one dam line from Topigs Norsvin. These data were collected between 1970 and 2020, and the traits were average daily gain during performance testing (ADGT), average daily gain throughout the lifetime (ADGL), backfat thickness, and loin depth. These traits are part of the breeding goals of each line. However, there was more emphasis on reproduction traits than on production traits in the dam line. Details on the amount of data utilized in this study are in Table 1. The data were recorded on originally preselected animals (i.e. the animals preselected by Topigs Norsvin), with the sire line being much more balanced than the dam line, in terms of proportions males and females with records per generation (ratio of males with records to females with records is about 50:50 in the sire line and about 20:80 in the dam line). We studied impact of genomic preselection (GPS) in the two lines separately, because the traits we studied had different weights in breeding goals of the two lines. Ancestors from the same line and year of birth with unknown parents were considered to be a separate base population in the pedigree. Each base population was fitted as a genetic group to account for genetic trend and differences in origin and selection history [16].

**Table 1.** Data utilized in subsequent ssGBLUP^a^ evaluations following each preselection scenario, after quality control

| Data in the subsequent ssGBLUP evaluation/Preselection scenario | With records on animals in the validation generation |  |  | Without records on animals in the validation generation |  |  |
| --- | --- | --- | --- | --- | --- | --- |
|  | Reference<sup>b</sup> | VGP<sup>c</sup> | MGP<sup>d</sup> | Reference<sup>b</sup> | VGP<sup>c</sup> | MGP<sup>d</sup> |
| *The sire line (number of validation animals per trait is ± 1383)* |  |  |  |  |  |  |
| Number of animals in the pedigree | 81,875 | 60,950 | 12,777 | 81,875 | 60,950 | 12,777 |
| Number of animals with record for at least one trait<sup>e</sup> | 75,129 | 54,217 | 6,065 | 52,846 | 52,846 | 4,694 |
| Number of animals with genotypes | 33,506 | 23,315 | 5,131 | 33,506 | 23,315 | 5,131 |
| Number of SNP genotyped | 20,550 | 20,963 | 20,926 | 20,550 | 20,963 | 20,926 |
| *The dam line (number of validation animals per trait is ± 2051)* |  |  |  |  |  |  |
| Number of animals in the pedigree | 160,426 | 124,031 | 33,485 | 160,426 | 124,031 | 33,485 |
| Number of animals with record for at least one trait | 139,403 | 103,018 | 12,514 | 100,710 | 100,710 | 10,206 |
| Number of animals with genotypes | 50,895 | 36,369 | 9,072 | 50,895 | 36,369 | 9,072 |
| Number of SNP genotyped | 19,199 | 19,256 | 20,647 | 19,199 | 19,256 | 20,647 |

<sup>a</sup> single-step genomic best linear unbiased prediction
<sup>b</sup> In the reference scenario, the subsequent ssGBLUP evaluation utilized the entire available data until the validation generation
<sup>c</sup> Validation generation preselection (VGP) scenario, in which all animals in the validation generation without progeny in the data were discarded
<sup>d</sup> Multi-generation preselection (MGP) scenario, in which all animals in the validation and training generations without progeny in the data were discarded
<sup>e</sup> About 87% and 70% of the animals in sire and dam lines respectively have records for all the four traits utilized in this study, and even bigger numbers have records for any two and three traits. We decided to keep any animal with record for at least one of the traits (92% and 87% of the animals in the sire and dam lines respectively) because every animal in the analyses would benefit from records on relatives and records of correlated traits (see Table S1), in addition to its own record on the primary trait.

### Training and validation generations

Per line, we split all animals into two groups, according to a cut-off birth date. Animals born before or on the cut-off birth date were used as training population, and animals born after the cut-off birth date were used as validation population. The cut-off date (to split the data into training and validation populations) was 31^st^ January, 2017 for the sire line, and 31^st^ December, 2015 for the dam line. Then from the validation population, animals that met the following requirements were selected as validation animals: 1) none of their parents were in the validation population, and 2) the animals had phenotyped progeny. The first requirement ensured that our validation animals were from only one generation, and the second requirement enabled comparing GEBV of the validation animals against their progeny yield deviations (PYD) [17]. Since records on validation animals were included in some of our subsequent evaluation scenarios (as will be seen later), we chose to use PYD as proxy for true breeding values (TBV) because PYD are estimated from phenotypes that were not included in the subsequent genetic evaluations.

### Genomic data and quality control

Our genomic data included genotypes of animals for about 21,000 SNP segregating in both lines, and distributed across the 18 autosomes in the pig genome. The SNP were genotyped using a custom SNP chip. We used Plink [18] for all quality control operations on our genomic data. Per GPS scenario (as described later) and per line, animals and SNPs with call rates less than 90% were removed, as well as SNPs that deviated from Hardy-Weinberg equilibrium (Hardy-Weinberg equilibrium exact test p value = 10^−15^), or had a minor allele frequency below 0.005. Table 1 contains the summary of the pedigree, genomic and phenotypic information utilized in the subsequent genetic evaluations following each GPS scenario.

### Computation of precorrected phenotypes

In our genetic evaluations, we used precorrected phenotypes (rather than raw phenotypes) as records. Animals of different lines were sometimes raised together, so they shared some fixed and non-genetic random effects. Because we studied impact of GPS within lines, it was necessary to correct phenotypes for all non-genetic effects before the data was divided into lines. Another motivation for using precorrected phenotypes is that after implementing our additional GPS scenarios (as described in detail in the next section), some classes of these non-genetic effects could be left with only one or a few animals. Then correcting for these effects would be less accurate compared to correcting for them before implementing our additional GPS scenarios. To compute precorrected phenotypes (**y**_**c**_), we first ran the following multi-trait pedigree-based animal model as follows:

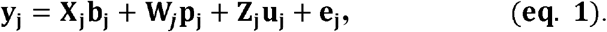

Where for every trait (j) **Y**_**j**_ was the vector of phenotypes; **b**_**j**_ was the vector of fixed effects, with incidence matrix **X**_**j**_ **; P**_**j**_ was the vector of non-genetic random effects, with incidence matrix **W**_**j**_**; u**_**j**_ was the vector of breeding values, with incidence matrix **z_j_** ; and **e**_**j**_ was the vector of residuals. The model assumed **u**_**j**_ and **e**_**j**_ to be normally distributed, each with mean of zero. For all traits (and across all animals), **u** and **e** had variance-covariance matrices **A** ⊗ **G** and **I** ⊗ **R**, respectively. Where **A** was the pedigree relationship matrix among animals, **I** was an identity matrix with dimensions equal to the number of animals with records, and **G** and **R** were respectively the trait by trait additive genetic and residual variance-covariance matrices. Then for every animal (i) with phenotype for trait j, we computed its precorrected phenotype (*Y*_*cij*_) as:

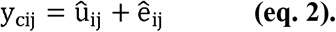

The (co)variance components used for this analysis were estimated, before separating the data into lines, from a four-trait pedigree-based animal model in ASReml [19] using **eq. 1**. All computations of (G)EBV were performed using MiXBLUP [20]. We decided to use a pedigree-based model (instead of a single-step model) to estimate the variance components because previous studies [21,22] showed that in populations undergoing genomic selection (as in our data), pedigree-based models estimate variance components in the pedigree founders at least as good as single-step models.

### Preselection

Per line, we implemented a reference scenario and two scenarios that added layers of GPS. The reference scenario - against which other scenarios could be compared -only included the original GPS implemented by Topigs Norsvin. Thus, the subsequent ssGBLUP evaluations following the reference scenario utilized the entire available data until the validation generation. The second scenario is called validation generation preselection (the VGP scenario). In this scenario, we only implemented additional GPS in the validation generation, by discarding all animals in the validation generation that had no progeny in the data, but had genotypes and/or phenotypes. The third scenario is called multi-generation preselection (the MGP scenario), in which we discarded any animal in the validation and training generations without progeny in the data. Animals kept after each of the GPS scenarios are shown in Figure 1.

**Figure 1.**
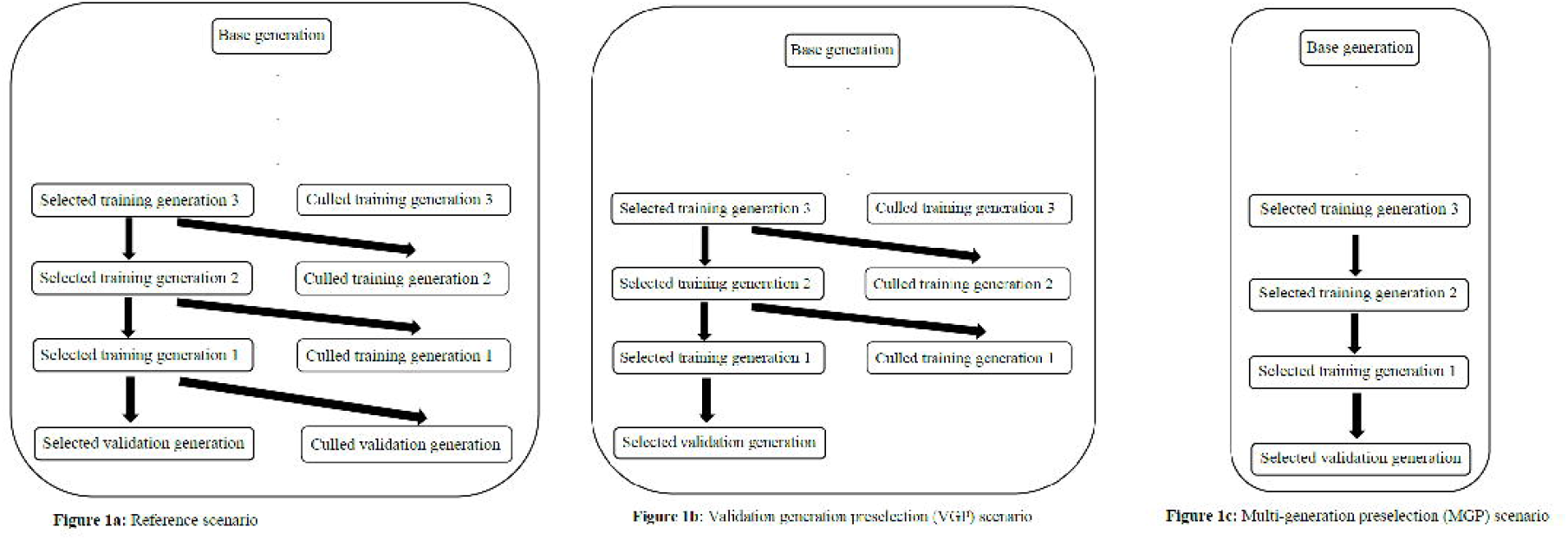
Overview of groups of animals used in subsequent ssGBLUP for each of the considered GPS scenarios

### Subsequent genetic evaluations

Following every scenario of GPS, we implemented a subsequent ssGBLUP evaluation with all animals that survived the GPS. We call this evaluation subsequent because it came after the initial evaluation that provided the GEBV used in preselection. The ssGBLUP evaluations were conducted with and without records (i.e. own precorrected phenotypes) on the animals in the validation generation (see Table 1), to represent traits with records (e,g, production traits) and those without records (e,g. reproduction traits) available during subsequent evaluations. Progeny of validation animals were not included in the subsequent genetic evaluations. We estimated variance components for every preselection scenario, per line, using a pedigree-based multi-trait animal model in ASReml. We used these scenario-specific variance components in the subsequent genetic evaluations to ensure that the variance components used were appropriate for the precorrected phenotypes. At the subsequent genetic evaluations, the (multi-trait) model used for the estimations of both variance components and breeding values for every trait (j) was:

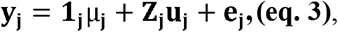

where for every trait (j) **y** _**j**_ was the vector of precorrected phenotypes; **1** _**j**_ was an incidence vector of 1’s, and was incidence matrix, linking precorrected phenotypes to overall mean and random animal effects, respectively; **μ**_**j**_ was the overall mean; **u**_**j**_ was the vector of breeding values; and was the vector of residuals. The model assumed **u** _**j**_ and to be normally distributed, each with mean of zero. For all traits (and across all animals), **u**_**j**_ and **e** _**j**_ had variance-covariance matrices **H** ⊗ **G** and **I** ⊗ **R**, respectively. Where **H** was the combined genomic and pedigree relationship matrix among animals as explained hereafter, **I** was an identity matrix with dimensions equal to the number of animals with records, and **G** and **R** were respectively the trait by trait additive genetic and residual variance-covariance matrices. We also repeated all subsequent genetic evaluations using PBLUP, to verify the impact of using genotypes on the observed results.

### Implementation of single-step GBLUP

The inverse of the combined pedigree-genomic relationship (**H**^-1^) was obtained as follows [23,24]:

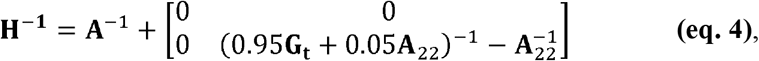

where **A**^-1^ was the inverse of the pedigree relationship matrix, and **A**_22_ was part of the pedigree relationship matrix referring to genotyped animals. We considered inbreeding in setting up both **A**^-1^ and **A**_22_, as ignoring inbreeding in setting up **A**^-1^ and **A**_22_ has been reported to cause bias in GEBV [13]. The adjusted genomic relationship matrix **G**_**t**_ was computed as follows [14,25]:

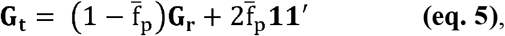

where 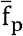 was the average pedigree inbreeding coefficient across genotyped animals, **G**_**t**_ was the raw genomic relationship matrix computed following the first method of VanRaden [26], and **11**^′^ was a matrix of 1s. As the animals with genotypes in this study were selectively genotyped, this transformation made sure that the impact of selective genotyping was taken care of and that **G** and **A**_**22**_ were on the same scale and therefore compatible [14,15]. To compute **G_r_**, we computed (current) allele frequencies using all available genomic data after quality control. We gave the weights of 0.95 to **G**_**t**_ and 0.05 to **A**_**22**_ to ensure that **G** was invertible [23,24].

### Measures of accuracy and bias in the subsequent genetic evaluations

We used progeny yield deviation (PYD) [17] as a proxy for true breeding value (TBV), against which GEBV were compared when computing accuracy and bias. To compute PYD, we ran a multi-trait pedigree-based animal model per line in MiXBLUP, with precorrected phenotypes as records and an overall mean as the only fixed effect **(eq. 3)**. The (co)variance components used in this model were also estimated per line in ASReml, from precorrected phenotypes, using a multi-trait pedigree-based animal model that only included a mean fixed effect **(eq. 3)**. From the output of this analysis, we computed PYD for all validation sires and dams (i) for each trait (j) as:

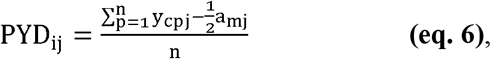

where PYD_ij_ was the progeny yield deviation of a sire or dam i for trait j, y_cpj_ was the precorrected phenotype of a progeny p of the sire or dam i for trait j, a_mj_ was the breeding value of the mate of sire or dam i (for trait j) in producing offspring p, and n was the number of phenotyped progeny of sire or dam i. Estimation of PYD was done before removing progeny of validation animals from the data. Since progeny of validation animals were not included in subsequent genetic evaluations, comparing (G)EBV to PYD can be considered as a forward-in-time validation. We computed approximate reliability of PYD for each validation animal for each trait, and used this approximate reliability as the weighting factor to compute accuracy and bias, to account for differences in number of progeny used to estimate PYD for different validation animals. The reliability of PYD was approximated as:

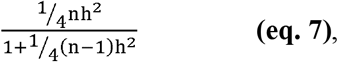

where *n* was the validation animal’s number of half-sib progeny with records, and *h*^*2*^ was the heritability of the trait [27]. For convenience, we assumed all progeny of a validation animal were half-sibs, though some of them were full-sibs. We also computed unweighted accuracy an bias, and accuracy and bias weighted by approximate PYD reliabilities obtained considering all progeny as full sibs. We did not observe statistically significant differences among the results, so we decided to only report the accuracy and bias weighted by approximate reliability computed considering all progeny as half sibs (as in eq. 7).

Validation accuracy was computed as weighted Pearson’s correlation coefficient between PYD and GEBV of all validation animals, using the ‘cor.test’ function of the ‘stats’ package in R [28]. We computed the standard errors (SE) of the estimates from the confidence intervals (CI) produced by the ‘cor.test’ function. Validation accuracy is not numerically the same as the accuracy of predicting TBV, since PYD has some non-genetic component, in addition to TBV [17]. However, validation accuracy and accuracy of predicting TBV increase and decrease together [29], and this property of validation accuracy enables us to use it to make comparison among subsequent genetic evaluation scenarios.

We computed two types of bias. The first type is level bias, which is a measure of whether estimated genetic gain is equal to true genetic gain. Level bias was computed as the weighted mean difference between PYD and half of the (G)EBV across all validation animals, expressed in additive genetic standard deviation (SD) units of the trait. We used the ‘weighted.mean’ function of the ‘stats’ package in R [28] to compute estimates of the weighted mean differences, and used the ‘weighted_se’ function of the ‘diagis’ package in R [30] to compute SE of the estimates. A negative difference means that GEBV are on average overestimated, and therefore genetic gain is overestimated, and vice versa. Since PYD were computed from a dataset that included information on progeny of validation animals and (G)EBV were computed without information on progeny of validation animals, PYD and (G)EBV were on different scales. Therefore before computing differences between PYD and half of the (G)EBV of validation animals, we scaled PYD and (G)EBV to be expressed against the same genetic base, consisting of the first three training generations. We did this in the following steps: from the model used in computing PYD, we computed average EBV across all animals in the first three training generations. We then subtracted half of this average EBV from PYD of each validation animal. Then for each subsequent genetic evaluation, we computed the average (G)EBV of all animals in the first three training generations. We then subtracted this average (G)EBV from (G)EBV of each validation animal.

The other type of bias we computed is dispersion bias, which was measured by the weighted regression coefficient of PYD on (G)EBV of all validation animals. We used the ‘lm’ function of the ‘stats’ package in R [28] to compute both the estimates and SE of the regression coefficients. If the regression coefficient is equal to the expected value, then there is no dispersion bias. Note that the expected value is 0.5, because PYD only includes half of the breeding value of a parent. A regression coefficient less than the expected value means that variance of (G)EBV is inflated, and vice versa.

## Results

Table 2 shows normalized means and SD of precorrected phenotypes of the traits analyzed, following the implememted genomic preselection (GPS) scenarios. The results show that our additionally implemented GPS was effective, as the lines were selected (and preselected) for increased feed efficiency (i.e. higher feed intake and higher average daily gain), slightly decreased backfat thickness, and slightly increased loin depth. As the validation generation for both validation generation preselection (VGP) and multi-generation preselection (MGP) scenarios only contained the preselected animals, means and SD of precorrected phenotypes of the traits are the same for these two GPS scenarios when only considering the animals in the validation generation. When only the validation generation was considered (i.e. the middle part of Table 2), means of precorrected phenotypes of average daily gain during performance testing (ADGT) and average daily gain throughout life (ADGL) increased from reference scenario to VGP and MGP scenarios. At the same time, SD of precorrected phenotypes of these traits decreased from reference scenario to VGP and MGP scenarios. In other words, means of precorrected phenotypes of these traits were higher, and SD of the precorrected phenotypes were lower, among preselected animals in the validation generation than among all animals in the validation generation. For backfat thickness, both mean and SD slightly decreased from reference to VGP and MGP scenarios. The change was also only slight for loin depth, with the mean slightly increasing, and the SD slightly decreasing, from reference to VGP and MGP scenarios. Higher effectiveness of MGP over VGP can be seen from the right part of Table 2 (i.e. when means and SD were computed across the entire data). For the positively (pre)selected traits (i.e ADGT, ADGL, and loin depth) means were higher for MGP than for VGP. For backfact, which was negatively (pre)selected, mean was lower for MGP than for VGP. As would be expected, SD were in all cases lower for MGP than for VGP.

**Table 2.** Normalized^a^ Means and SD (in brackets) of precorrected phenotypes of the traits utilized in this study, following each GPS scenario

| Trait /Preselection scenario | Only within the validation generation |  | Across the entire data |  |
| --- | --- | --- | --- | --- |
|  | Reference<sup>b</sup> | VGP<sup>c</sup>/MGP<sup>d</sup> | VGP | MGP |
| *Sire line* |  |  |  |  |
| ADGT<sup>e</sup> (g/day) | 0.12 (1.00) | 0.51 (0.80) | -0.08 (1.00) | 0.38 (0.84) |
| ADGL<sup>f</sup> (g/day) | 0.03 (1.00) | 0.41 (0.85) | -0.11 (1.00) | 0.31 (0.86) |
| Backfat thickness (mm) | -0.27 (1.00) | -0.29 (0.95) | -0.01 (1.00) | -0.10 (0.96) |
| Loin depth (mm) | 0.26 (1.00) | 0.26 (0.97) | 0.17 (1.00) | 0.20 (0.97) |
| *Dam line* |  |  |  |  |
| ADGT (g/day) | 0.22 (1.00) | 0.62 (0.90) | -0.06 (1.00) | 0.40 (0.91) |
| ADGL (g/day) | 0.11 (1.00) | 0.53 (0.87) | -0.11 (1.00) | 0.32 (0.90) |
| Backfat thickness (mm) | -0.13 (1.00) | 0.05 (0.97) | -0.02 (1.00) | -0.06 (0.99) |
| Loin depth (mm) | 0.22 (1.00) | 0.13 (0.98) | 0.20 (1.00) | 0.24 (0.99) |

<sup>a</sup> The values were normalized by dividing them by the standard deviations of their corresponding reference scenarios
<sup>b</sup> In the reference scenario, the subsequent ssGBLUP evaluation utilized the entire available data until the validation generation
<sup>c</sup> Validation generation preselection (VGP) scenario, in which all animals in the validation generation without progeny in the data were discarded
<sup>d</sup> Multi-generation preselection (MGP) scenario, in which all animals in the validation and training generations without progeny in the data were discarded
<sup>e</sup> ADGT: Average daily gain during performance testing
<sup>f</sup> ADGL: ADG throughout life

Results of the subsequent genetic evaluations conducted with ssGBLUP are presented in Tables 3 and 4, respectively for the sire line and the dam line. Results in Tables 5 and 6 are from subsequent genetic evaluations done with PBLUP, respectively for the sire line and the dam line. For every parameter in these tables (i.e. estimated heritability, validation accuracy, level bias, and dispersion bias), we showed the estimate and SE of the estimate. We always used a one-tailed two-sample t-test at 5% significance level to determine whether two estimates were different. We included the estimated heritabilities in our results because they help in explaining the results of accuracy and bias.

**Table 3.**
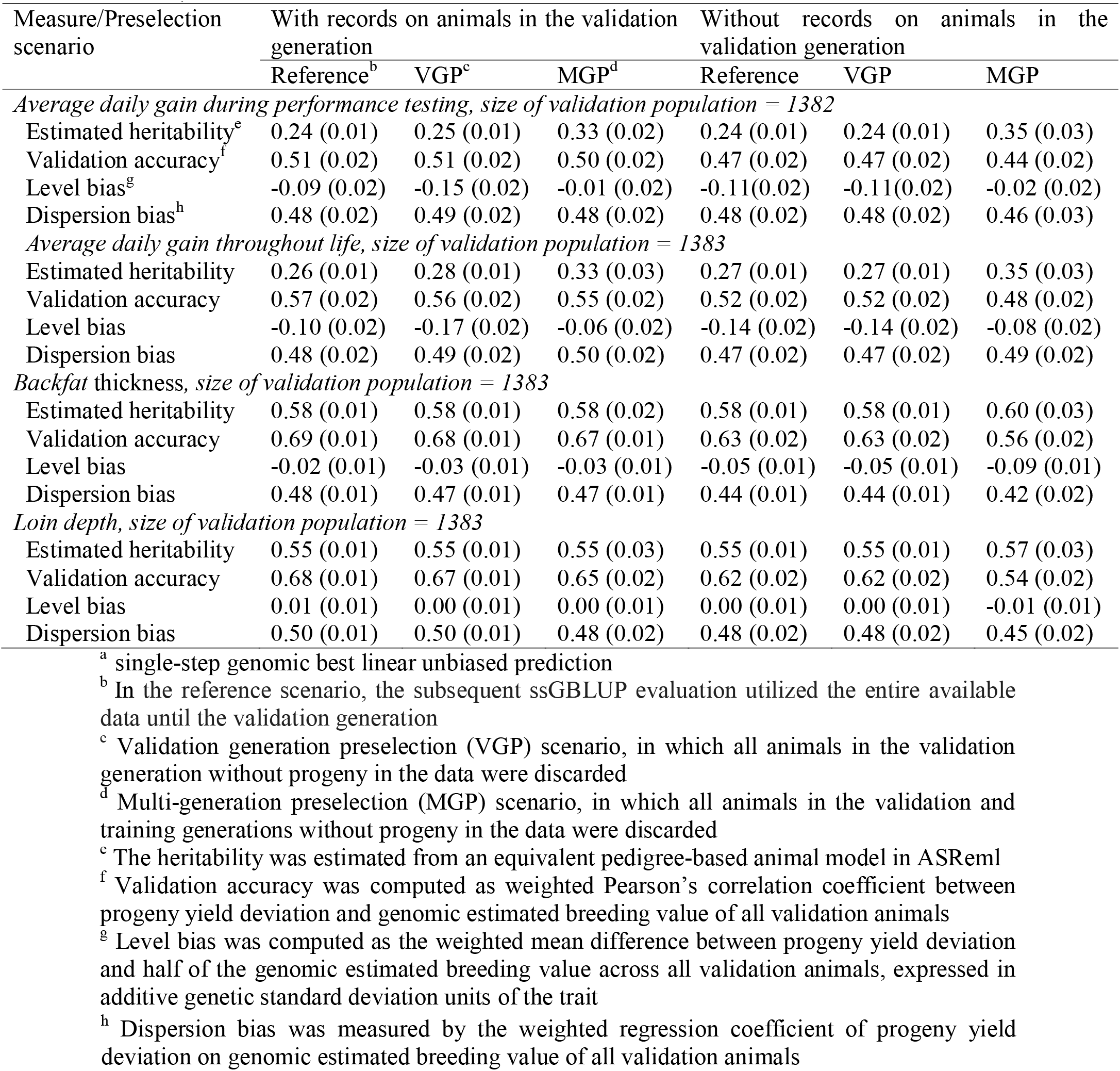
Performance of ssGBLUP^a^ in the subsequent evaluations in the sire line (SE in brackets)

**Table 4.**
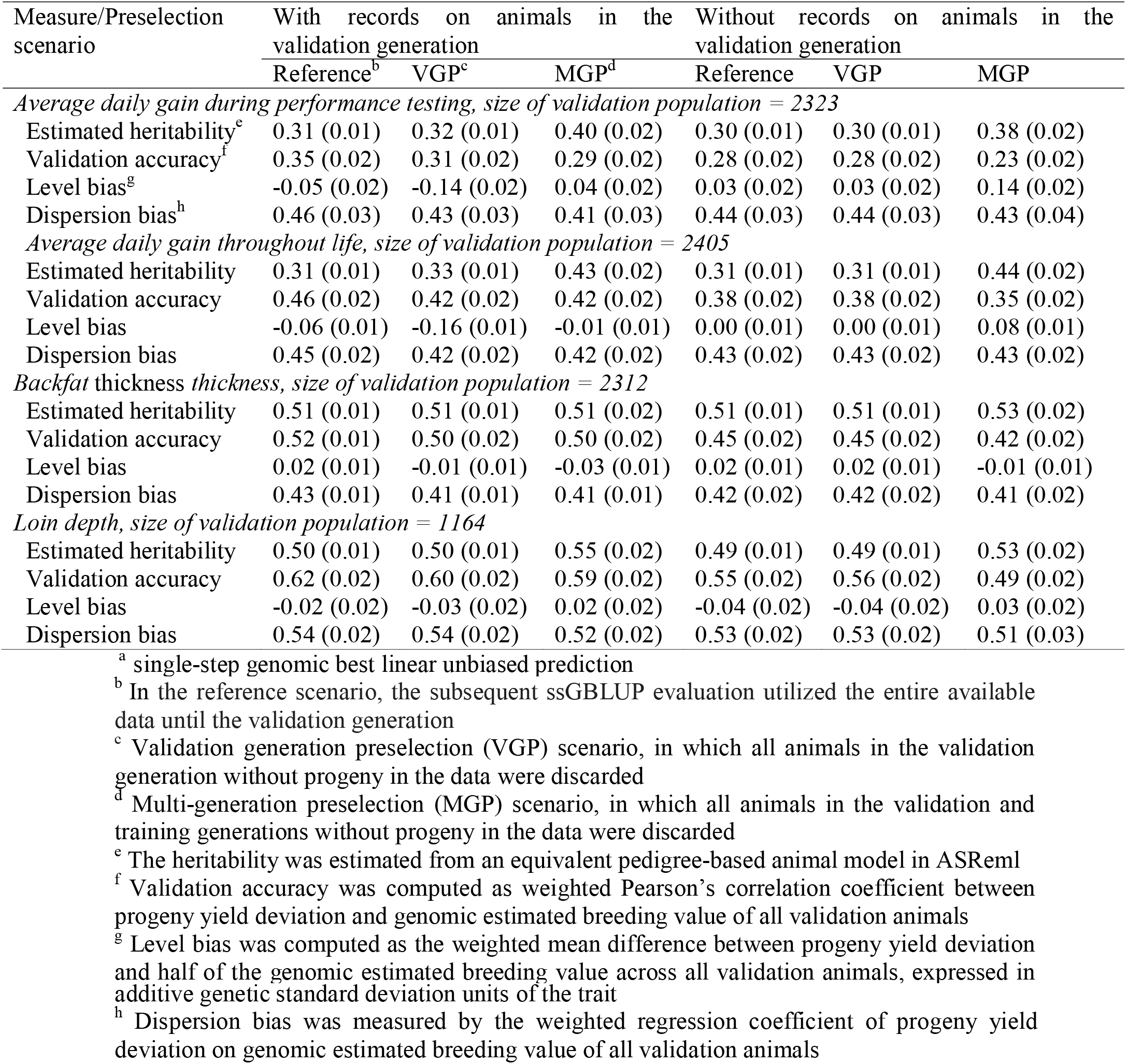
Performance of ssGBLUP^a^ in the subsequent evaluations in the dam line (SE in brackets)

| Measure/Preselection scenario | With records on animals in the validation generation |  |  | Without records on animals in the validation generation |  |  |
| --- | --- | --- | --- | --- | --- | --- |
|  | Reference<sup>b</sup> | VGP<sup>c</sup> | MGP<sup>d</sup> | Reference | VGP | MGP |
| *Average daily gain during performance testing, size of validation population = 2323* |  |  |  |  |  |  |
| Estimated heritability<sup>e</sup> | 0.31 (0.01) | 0.32 (0.01) | 0.40 (0.02) | 0.30 (0.01) | 0.30 (0.01) | 0.38 (0.02) |
| Validation accuracy<sup>f</sup> | 0.35 (0.02) | 0.31 (0.02) | 0.29 (0.02) | 0.28 (0.02) | 0.28 (0.02) | 0.23 (0.02) |
| Level bias<sup>g</sup> | -0.05 (0.02) | -0.14 (0.02) | 0.04 (0.02) | 0.03 (0.02) | 0.03 (0.02) | 0.14 (0.02) |
| Dispersion bias<sup>h</sup> | 0.46 (0.03) | 0.43 (0.03) | 0.41 (0.03) | 0.44 (0.03) | 0.44 (0.03) | 0.43 (0.04) |
| *Average daily gain throughout life, size of validation population = 2405* |  |  |  |  |  |  |
| Estimated heritability | 0.31 (0.01) | 0.33 (0.01) | 0.43 (0.02) | 0.31 (0.01) | 0.31 (0.01) | 0.44 (0.02) |
| Validation accuracy | 0.46 (0.02) | 0.42 (0.02) | 0.42 (0.02) | 0.38 (0.02) | 0.38 (0.02) | 0.35 (0.02) |
| Level bias | -0.06 (0.01) | -0.16 (0.01) | -0.01 (0.01) | 0.00 (0.01) | 0.00 (0.01) | 0.08 (0.01) |
| Dispersion bias | 0.45 (0.02) | 0.42 (0.02) | 0.42 (0.02) | 0.43 (0.02) | 0.43 (0.02) | 0.43 (0.02) |
| *Backfat thickness thickness, size of validation population = 2312* |  |  |  |  |  |  |
| Estimated heritability | 0.51 (0.01) | 0.51 (0.01) | 0.51 (0.02) | 0.51 (0.01) | 0.51 (0.01) | 0.53 (0.02) |
| Validation accuracy | 0.52 (0.01) | 0.50 (0.02) | 0.50 (0.02) | 0.45 (0.02) | 0.45 (0.02) | 0.42 (0.02) |
| Level bias | 0.02 (0.01) | -0.01 (0.01) | -0.03 (0.01) | 0.02 (0.01) | 0.02 (0.01) | -0.01 (0.01) |
| Dispersion bias | 0.43 (0.01) | 0.41 (0.01) | 0.41 (0.01) | 0.42 (0.02) | 0.42 (0.02) | 0.41 (0.02) |
| *Loin depth, size of validation population = 1164* |  |  |  |  |  |  |
| Estimated heritability | 0.50 (0.01) | 0.50 (0.01) | 0.55 (0.02) | 0.49 (0.01) | 0.49 (0.01) | 0.53 (0.02) |
| Validation accuracy | 0.62 (0.02) | 0.60 (0.02) | 0.59 (0.02) | 0.55 (0.02) | 0.56 (0.02) | 0.49 (0.02) |
| Level bias | -0.02 (0.02) | -0.03 (0.02) | 0.02 (0.02) | -0.04 (0.02) | -0.04 (0.02) | 0.03 (0.02) |
| Dispersion bias | 0.54 (0.02) | 0.54 (0.02) | 0.52 (0.02) | 0.53 (0.02) | 0.53 (0.02) | 0.51 (0.03) |

<sup>a</sup> single-step genomic best linear unbiased prediction
<sup>b</sup> In the reference scenario, the subsequent ssGBLUP evaluation utilized the entire available data until the validation generation
<sup>c</sup> Validation generation preselection (VGP) scenario, in which all animals in the validation generation without progeny in the data were discarded
<sup>d</sup> Multi-generation preselection (MGP) scenario, in which all animals in the validation and training generations without progeny in the data were discarded
<sup>e</sup> The heritability was estimated from an equivalent pedigree-based animal model in ASReml
<sup>f</sup> Validation accuracy was computed as weighted Pearson's correlation coefficient between progeny yield deviation and genomic estimated breeding value of all validation animals
<sup>g</sup> Level bias was computed as the weighted mean difference between progeny yield deviation and half of the genomic estimated breeding value across all validation animals, expressed in
additive genetic standard deviation units of the trait <sup>h</sup> Dispersion bias was measured by the weighted regression coefficient of progeny yield deviation on genomic estimated breeding value of all validation animals

**Table 5.** Performance of PBLUP^a^ in the subsequent evaluations in the sire line (SE in brackets)

| Measure/Preselection scenario | With records on animals in the validation generation |  |  | Without records on animals in the validation generation |  |  |
| --- | --- | --- | --- | --- | --- | --- |
|  | Reference<sup>b</sup> | VGP<sup>c</sup> | MGP<sup>d</sup> | Reference | VGP | MGP |
| *Average daily gain during performance testing, size of validation population = 1382* |  |  |  |  |  |  |
| Estimated heritability | 0.24 (0.01) | 0.25 (0.01) | 0.33 (0.02) | 0.24 (0.01) | 0.24 (0.01) | 0.35 (0.03) |
| Validation accuracy<sup>e</sup> | 0.51 (0.02) | 0.50 (0.02) | 0.49 (0.02) | 0.41 (0.02) | 0.41 (0.02) | 0.40 (0.02) |
| Level bias<sup>f</sup> | -0.04 (0.02) | -0.11 (0.02) | 0.01 (0.02) | -0.01 (0.02) | -0.01 (0.02) | 0.01 (0.02) |
| Dispersion bias<sup>g</sup> | 0.53 (0.02) | 0.54 (0.03) | 0.48 (0.02) | 0.55 (0.03) | 0.55 (0.03) | 0.49 (0.03) |
| *Average daily gain throughout life, size of validation population = 1383* |  |  |  |  |  |  |
| Estimated heritability | 0.26 (0.01) | 0.28 (0.01) | 0.33 (0.03) | 0.27 (0.01) | 0.27 (0.01) | 0.35 (0.03) |
| Validation accuracy | 0.58 (0.02) | 0.56 (0.02) | 0.54 (0.02) | 0.47 (0.02) | 0.47 (0.02) | 0.44 (0.02) |
| Level bias | -0.06 (0.02) | -0.14 (0.02) | -0.04 (0.02) | -0.05 (0.02) | -0.05 (0.02) | -0.05 (0.02) |
| Dispersion bias | 0.55 (0.02) | 0.55 (0.02) | 0.51 (0.02) | 0.56 (0.03) | 0.56 (0.03) | 0.54 (0.03) |
| *Backfat thickness thickness, size of validation population = 1383* |  |  |  |  |  |  |
| Estimated heritability | 0.58 (0.01) | 0.58 (0.01) | 0.58 (0.02) | 0.58 (0.01) | 0.58 (0.01) | 0.60 (0.03) |
| Validation accuracy | 0.67 (0.01) | 0.66 (0.02) | 0.66 (0.02) | 0.48 (0.02) | 0.48 (0.02) | 0.46 (0.02) |
| Level bias | -0.03 (0.01) | -0.03 (0.01) | -0.03 (0.01) | -0.09 (0.01) | -0.09 (0.01) | -0.10 (0.01) |
| Dispersion bias | 0.50 (0.01) | 0.50 (0.02) | 0.50 (0.02) | 0.46 (0.02) | 0.46 (0.02) | 0.43 (0.02) |
| *Loin depth, size of validation population = 1383* |  |  |  |  |  |  |
| Estimated heritability | 0.55 (0.01) | 0.55 (0.01) | 0.55 (0.03) | 0.55 (0.01) | 0.55 (0.01) | 0.57 (0.03) |
| Validation accuracy | 0.66 (0.02) | 0.65 (0.02) | 0.64 (0.02) | 0.49 (0.02) | 0.49 (0.02) | 0.46 (0.02) |
| Level bias | 0.00 (0.01) | 0.00 (0.01) | 0.00 (0.01) | 0.01 (0.01) | 0.01 (0.01) | 0.00 (0.01) |
| Dispersion bias | 0.50 (0.02) | 0.49 (0.02) | 0.49 (0.02) | 0.48 (0.02) | 0.48 (0.02) | 0.46 (0.02) |

<sup>a</sup> Pedigree-based best linear unbiased prediction
<sup>b</sup> In the reference scenario, the subsequent PBLUP evaluation utilized the entire available data until the validation generation
<sup>c</sup> Validation generation preselection (VGP) scenario, in which all animals in the validation generation without progeny in the data were discarded
<sup>d</sup> Multi-generation preselection (MGP) scenario, in which all animals in the validation and training generations without progeny in the data were discarded
<sup>e</sup> Validation accuracy was computed as weighted Pearson's correlation coefficient between progeny yield deviation and genomic estimated breeding value of all validation animals
<sup>f</sup> Level bias was computed as the weighted mean difference between progeny yield deviation and half of the genomic estimated breeding value across all validation animals, expressed in additive genetic standard deviation units of the trait
<sup>g</sup> Dispersion bias was measured by the weighted regression coefficient of progeny yield deviation on genomic estimated breeding value of all validation animals

**Table 6.**
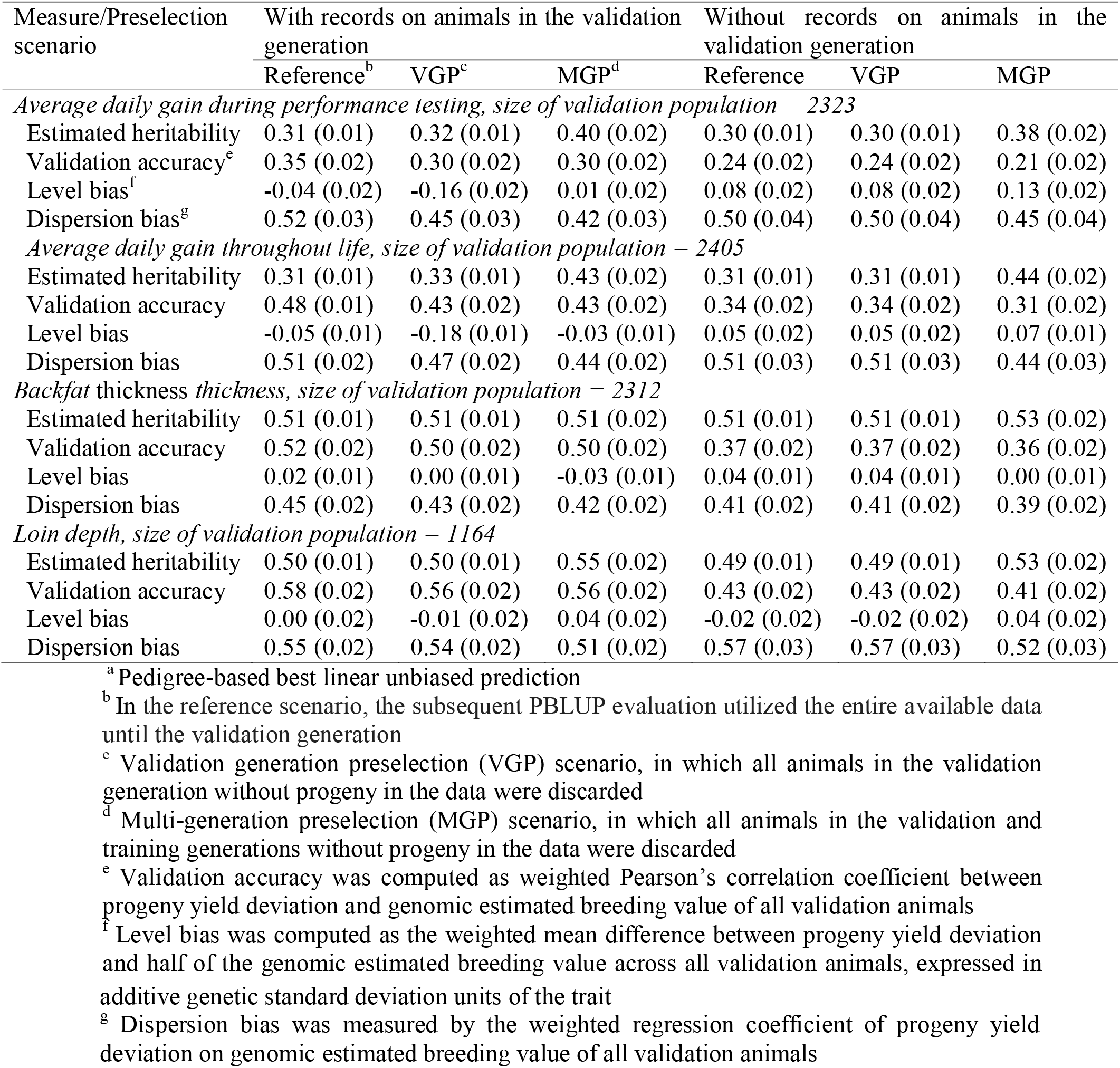
Performance of PBLUP^a^ in the subsequent evaluations in the dam line (SE in brackets)

Estimated heritabilities for ADGT and ADGL did not differ between reference and VGP scenarios, but increased in MGP scenarios. This was observed in both lines. For backfat thickness, heritabilities did not differ across GPS scenarios, neither in the sire line nor in the dam line. For loin depth, the heritabilities did not differ across GPS scenarios in the sire line. In the dam line however, the heritability of loin depth was higher in MGP scenarios than in reference and VGP scenarios. The above trends in heritabilities for all traits were observed whether records on animals in the validation generation were included or excluded in estimating the heritabilities. Without records on animals in the validation generation in the subsequent ssGBLUP evaluations, all results (including estimated heritabilities) for reference and VGP scenarios were the same. The increases in heritabilities observed with more preselection were generally due to decreases in residual variances with more preselection, while additive genetic variances generally did not differ across GPS scenarios (Tables S2 and S3).

### Subsequent ssGBLUP evaluations with records on animals in the validation generation

Validation accuracies did not differ across GPS scenarios for all traits in both lines, except for ADGT in the dam line, where the accuracy was lower in MGP scenario than in reference scenario (Table 4). Tendencies (i.e. indications that may not be statistically significant) towards lower accuracies with more GPS were however observed for all traits in both lines (Tables 3 and 4). In both lines, level bias was absent in all scenarios for loin depth, and only in some scenarios for ADGT, ADGL and backfat thickness. Even when level bias was present, it was still only marginal. The highest value of level bias recorded was -0.17 additive genetic SD units, under the VGP scenario for ADGL in the sire line (Table 3). Dispersion bias was absent (i.e. the regression coefficient of PYD on GEBV did not differ from its expected value of 0.5) for all traits in the sire line (Table 3), except in VGP and MGP scenarios for backfat thickness, where there was inflation (i.e. the regression coefficient was less than 0.5). On the other hand, dispersion bias was present for all traits in the dam line (Table 4), except in the reference scenario of ADGT and MGP scenario of loin depth. In the instances with dispersion bias in the dam line, the regression coefficients were greater than 0.5 (i.e. they were deflated) in reference and VGP scenarios of loin depth, and less than 0.5 in all other instances. However, the estimates of the regression coefficients did not differ across GPS scenarios within traits and lines (Tables 3 and 4), although they showed tendencies to decrease from reference to VGP to MGP scenarios.

### Subsequent ssGBLUP evaluations without records on animals in the validation generation

For ADGT, validation accuracy did not differ across GPS scenarios in the sire line (Table 3), but in the dam line it was lower in MGP scenario than in reference/VGP scenarios (Table 4). For ADGL, validation accuracies did not differ across GPS scenarios in both lines, but only tended to decrease from reference/VGP to MGP scenarios (Tables 3 and 4). For backfat thickness, validation accuracy in the sire line decreased from reference/VGP scenarios to MGP scenario, but did not differ across GPS scenarios in the dam line. For loin depth, validation accuracies in both lines decreased from reference/VGP to MGP scenarios. Level bias was present in reference and VGP scenarios for ADGT in the sire line, but absent in MGP scenario. The reverse is the case in the dam line, where level bias was absent in reference and VGP scenarios for ADGT, and present in MGP scenario. For ADGL, level bias was present in all GPS scenarios in the sire line, and in the dam only present in the MGP scenario. For backfat thickness, level bias was present across all GPS scenarios in the sire line, and absent across all GPS scenarios in the dam line. For loin depth, level bias was absent across all GPS scenarios in both lines. Although level bias was present in many scenarios, it was still only marginal, with ±0.14 additive genetic SD units being its highest estimate (Tables 3 and 4). For ADGT and ADGL, dispersion bias was absent across GPS scenarios in the sire line, and present across GPS scenarios in the dam line. For backfat thickness, dispersion bias was present across all scenarios in both lines. For loin depth, dispersion bias was absent across all scenarios in both lines, except in the MGP scenario in the sire line, where there was inflation. Here too, just as when records on animals in the validation generation were included in the subsequent evaluations, the estimates of the regression coefficients did not differ across all GPS scenarios within traits, and this was observed for all traits in both lines.

### Subsequent genetic evaluations with PBLUP

When records on animals in the validation generation were included in the subsequent evaluations, corresponding validation accuracies did not differ between PBLUP and ssGBLUP. This was observed for all traits in both lines. However, when records on animals in the validation generation were excluded from the subsequent evaluations in the sire line, validation accuracies of all traits were lower with PBLUP than with ssGBLUP. When records on animals in the validation generation were excluded from the subsequent evaluations in the dam line, validation accuracies did not differ between PBLUP and ssGBLUP for ADGT and ADGL, but were lower with PBLUP than with ssGBLUP for backfat thickness and loin depth. Just as validation accuracies from subsequent ssGBLUP evaluations, validation accuracies from subsequent PBLUP evaluations in most instances did not differ across corresponding GPS scenarios. Just like when the subsequent evaluations were done with ssGBLUP, here too, level bias with PBLUP in most cases did not differ from its corresponding value with ssGBLUP, and in many instances it was not different from zero. The highest value of level bias when the subsequent evaluations were done with PBLUP was -0.18 additive genetic SD units (i.e. in the VGP scenario for ADGL in the dam line when records on animals in the validation included were included in the subsequent evaluation; Table 6). Regression coefficients of PYD on (G)EBV in most instances did not differ between PBLUP and ssGBLUP, or from their expected value of 0.5. However, with PBLUP, the regression coefficients were sometimes bigger than 0.5 (e.g. in reference and VGP scenarios for ADGL in the sire line, and in reference and VGP scenarios for loin depth in the dam line). The regression coefficients were also in some instances bigger with PBLUP than with ssGBLUP (e.g. in reference and VGP scenarios for ADGT in the sire line, and in reference and VGP scenarios for ADGL in both lines). The regression coefficients with PBLUP were lower in MGP scenarios than in reference scenarios for ADGT and ADGL in the dam line when records on animals in the validation generation was included in the subsequent evaluations (Table 6). This is unlike with ssGBLUP, where the regression coefficients in all instances did not differ across corresponding GPS scenarios (Tables 3 and 4).

## Discussion

In this study, we investigated the impact of genomic preselection (GPS) on accuracy and bias in subsequent ssGBLUP evaluations of preselected animals. We used data from a commercial pig breeding program in which preselection has taken place, and retrospectively implemented additional layers of GPS. The data was on production traits of pigs from one sire line and one dam line. Per line, we implemented three GPS scenarios. We used precorrected phenotypes as records in the subsequent genetic evaluations, and progeny yield deviation (PYD) as the proxy for TBV. We did the subsequent genetic evaluations either with or without records on animals in the validation generation, and in all cases without progeny of validation animals. Validation accuracy decreased, or at least tended to, with more GPS. Dispersion bias was largely absent, and the regression coefficient of PYD on GEBV - the indicator of dispersion bias – in all instances did not differ among corresponding GPS scenarios. Level bias was also largely absent, and mean PYD minus Mean GEBV - the indicator of level bias - in most instances did not differ across GPS scenarios also. The above results were observed in both lines, for all traits, and whether records on animals in the validation generation were included in or excluded from the subsequent ssGBLUP evaluations.

Empirically assessing the impact of preselection on subsequent genetic evaluations of preselected animals requires comparison of scenarios taking into account different approaches, including scenarios without preselection. Since some GPS had already taken place in the dataset we used for this study, it was not possible to have a scenario without preselection. We therefore needed to come up with another way of investigating whether ssGBLUP is able to estimate GEBV in the subsequent evaluation of preselected animals without preselection bias in our current dataset and by extension in real breeding programs. We hypothesized that if ssGBLUP in subsequent evaluations yields unbiased GEBV for preselected animals despite additional GPS in the current dataset (i.e. in our VGP and MGP scenarios), then it also yields unbiased GEBV for preselected animals in the subsequent evaluations with the current dataset (i.e. in our reference scenarios). This is why we implemented the VGP and MGP scenarios, to implement additional layers of GPS over the regular GPS already implemented by the commercial pig breeding program. While the equivalent of VGP and MGP do not happen in real breeding programs, implementing these GPS scenarios in this study enabled us to investigate the impact of GPS on subsequent genetic evaluations of preselected animals using real data, by including different amounts of pedigree, genomic and phenotypic information in the subsequent genetic evaluations. Our results have shown that ssGBLUP in subsequent evaluations of pigs is indeed able to estimate GEBV of preselected animals without preselection bias. We believe that the findings of the current study can be extended to other animal breeding programs. This is because preselection, irrespective of its type and intensity and in how many stages it is implemented, has similar effects across breeding programs (i.e. preselection ensures that only better-than-average animals are phenotyped for the traits measured at advanced stages of lives of animals). The findings of the current study are in line with what we showed in our previous studies using simulated datasets [3,4], that in subsequent evaluations ssGBLUP estimates unbiased GEBV for preselected animals. In [3], ssGBLUP in subsequent evaluations estimated GEBV of preselected animals with accuracy loss compared to scenarios without preselection. We attributed the accuracy loss in scenarios with preselection to less numbers of sibs with records compared to the scenarios without preselection. In this study too, accuracy decreased, or at least tended to, with more preselection. This decrease can also be attributed to less numbers of relatives with records compared to the in scenarios with less preselection, as can be seen in Table 1.

The preselection we implemented in this study as well as in [3] and [4] are forms of non-ignorable selection (e.g. [9,31–33]). In the absence of genomic information, all the information utilized at these preselection stages need to be included in subsequent evaluations in order to avoid preselection bias (e.g. [9,12,34,35]). In our previous studies [3,4], we showed that ssGBLUP in subsequent evaluations of preselected animals estimates GEBV of preselected animals without preselection bias even if genotypes of preculled animals are not included. In [4], we also showed that genotypes of preculled animals are only needed in the subsequent ssGBLUP evaluations of their preselected sibs if their parents are not genotyped. Still in [4], we suggested that ssGBLUP uses genotypes of preselected animals and their parents to estimate the on-average positive Mendelian sampling terms of the preselected animals, and this enables ssGBLUP in subsequent evaluations to estimate GEBV of preselected animals without preselection bias. In this study preselected animals and their parents were genotyped, and indeed we did not observe preselection bias in our subsequent ssGBLUP evaluations despite not including genotypes of preculled animals in the subsequent evaluations.

### Comparison of results across preselection scenarios and between ssGBLUP and PBLUP

We have shown that for all traits, and especially for ADGT and ADGL, estimated heritabilities increased or at least tended to, with more GPS. These tendencies of the heritabilities of these traits to increase with more preselection were because changes in residual variances were bigger than their corresponding changes in additive genetic variances moving from reference to VGP to MGP scenarios (Tables S2 and S3). The likely explanation for this observation is that animals may have low phenotypes for non-genetic reasons such as injury, social stress or illness. This ‘dilutes’ the heritability. Typically, such poor-performing animals are not selected, and this results to selected groups of animals being more homogeneous in expressing their genetic potentials than unselected groups. This suggests heterogeneity of residual variances across herd-year classes for all traits, with the greatest heterogeneity in the reference scenarios. In both lines, we repeated the subsequent evaluations of the reference scenarios with records on animals in the validation generations included, with a model that corrects for heterogenous residual variances (results not shown; [20]). Since we found that estimates of accuracy and bias in these scenarios did not differ between our two models, we decided to continue with the simpler model (i.e. the model in eq. 3).

We have also shown that with both ssGBLUP and PBLUP, validation accuracy decreased or at least tended to, with more GPS. These tendencies can be explained by the fact that the amount of phenotypic information also reduced in that order (Table 1). The observed tendencies of heritabilities to increase with more preselection could have influenced, at least partly, the magnitudes of decreases in accuracies with decreases in amounts of phenotypic information due to preselection. This can contribute to explaining why decreases in validation accuracies with more GPS were in most instances not statistically significant. Although there were always tendencies that corresponding validation accuracies were higher with ssGBLUP than with PBLUP, we observed that the differences were in most instances not statistically significant. The fact that heritabilities were all relatively high (ranging from 0.24 to 0.58, Tables 3 to 6) explains, at least partly, the absence of significant differences between ssGBLUP and PBLUP evaluations when records on animals in the validation generation were included in the subsequent genetic evaluations. It is a common knowledge that the higher the heritability, the higher the importance of own performance information and the lesser the importance of genomic information in genetic evaluations (e.g. [17]).

We have shown that in this study, level bias was absent in most instances, and even when it was present it was only marginal. We have also shown that the measure of level bias (i.e. the difference between mean PYD and mean (G)EBV among validation animals) in most instances did not differ across corresponding GPS scenarios, regardless of whether ssGBLUP or PBLUP were used in the subsequent evaluations. In our previous study [3], we observed no level bias when ssGBLUP was used in subsequent genetic evaluations, irrespective of type or intensity of preselection. However, in [3], we found level bias to be increasing with intensity of preselection when we used PBLUP in subsequent genetic evaluations. Patry et al [1,10,11] also reported significant level bias when subsequent genetic evaluations of genomically preselected animals were done with PBLUP, except when some pseudo-phenotypic information on preculled animals was included in the subsequent PBLUP evaluations. Just like with level bias, we have shown that in this study dispersion bias was absent in most instances, regardless of whether ssGBLUP or PBLUP was used in the subsequent evaluations. In our previous study with a simulated dataset [3], we found that regression coefficients of TBV on (G)EBV - the indicator of dispersion bias - were bigger and closer to the expected value of 1 when ssGBLUP was used in the subsequent genetic evaluations compared to when PBLUP was used. In [3], we also found that the regression coefficient became smaller as preselection intensity increased when PBLUP was used, but did not differ when ssGBLUP was used, irrespective of preselection intensity. Preselection and subsequent selection were multi-trait in this study, and single trait in the previous studies [1,3,10,11]. This means that the chance of having multiple litter mates left in the data after preselection and subsequent selection is higher in the current study than in the previous studies. With multiple litter mates with records in the data, MS terms can be estimated reasonably well even in the absence of genomic information. This is likely the reason we did not observe level or dispersion bias in subsequent PBLUP evaluations of preselected animals in this study.

In the absence of selection, the expectation of regression coefficient of PYD on (G)EBV is 0.5, because PYD only represents half of the breeding value of the parent. However, when validation animals are on average better than the average of their age group, the expectation of the regression coefficient decreases in single-trait subsequent evaluations, depending on how much the validation animals deviate from the average of their age group (e.g. [36,37]). In the data used in this study, ADGT and ADGL had heavier weights in the breeding goals of the two lines than backfat thickness and loin depth, so we expected that our GPS would have smaller impacts on the regression coefficients for backfat thickness and loin depth than for ADGT and ADGL. We however did not observe smaller regression coefficients or regression coefficients that were further away from 0.5 for ADGT and ADGL than for backfat thickness and loin depth, neither with ssGBLUP nor with PBLUP. As explained in the previous paragraph, the fact that we implemented multi-trait subsequent evaluations in this study, as opposed to the single-trait subsequent evaluations in [36] and [37], could explain the differences between these two groups of studies.

### Comparison of results across the two lines

Even in the dam line where the original GPS was at least numerically more intense and ratio of males with records to females with records in any generation was about 20:80, we generally did not observe significant decrease in validation accuracy, or increase in level and dispersion biases across our GPS scenarios. We however found that corresponding validation accuracies for ADGT, ADGL and backfat thickness were always higher in the sire line than in the dam line, despite the corresponding estimated heritabilities for ADGT and ADGL in many instances being higher in the dam line than in the sire line. These higher accuracies in the sire line than in the dam line may be explained by the relatively higher phenotyping and genotyping rates in the sire line than in the dam line (Table 1), meaning that validation animals in the sire line had more relatives with phenotypes and/or genotypes than validation animals in the dam lines.

### Genotypes of preculled animals did not affect the subsequent ssGBLUP evaluations

In the subsequent ssGBLUP evaluations without records on animals in the validation generation, results from corresponding reference and VGP scenarios are exactly the same, at least up to two decimal places (Tables 3 and 4). However, in terms of data content, reference scenarios contained genotypes of the animals preculled in the corresponding VGP scenarios, in addition to all the data contained in the corresponding VGP scenarios (Table 1). The fact that results from these two scenarios are the same means that genotypes of the preculled animals did not affect the reference scenarios. In this study, most (about 95%) of the validation animals and their parents had genotypes. This is in line with the conclusion from our previous study [4], that genotypes of preculled animals are only useful in subsequent ssGBLUP evaluations of their preselected sibs when their parents are not genotyped.

### Potential additional sources of bias in ssGBLUP from our data

In practical datasets as used in this study, it is difficult to completely rule out some mistakes in pedigree recording and in genotyping. At our genomic data quality control stage, genotypes of a few thousand animals were discarded because the animals did not meet the genomic data quality standard (of being genotyped for at least 90% of the SNP). Genotyping mistakes could still not be completely ruled out in the genomic data that passed quality control. In Tables 3 to 6, we saw that for some traits, heritabilities were different across the implemented GPS scenarios, although the animals in the base generation were the same. This implies that different subsets of the same data gave rise to different estimated (co)variance components in the base generation, and that it is likely that after some of the GPS scenarios were implemented, the estimated (co)variance components were different from their true values, at least for some of the traits. While these are all potential additional sources of bias in ssGBLUP evaluations, they are difficult to avoid in practice [13]. However, in general, we have shown that these potential additional sources of bias did not cause significant bias in our ssGBLUP evaluations, as both level and dispersion biases were in most cases absent, and even when present they were only marginal and mostly not different across corresponding GPS scenarios.

## Conclusions

When subsequent genetic evaluations of preselected animals are done with ssGBLUP, genomic preselection in single or multiple generations only slightly decreases realized accuracy, and hardly causes level or dispersion bias. This conclusion is expected to hold regardless of whether records on animals in the validation generation are included or excluded in the subsequent evaluations, and regardless of the weight of the trait in the breeding goal. Although these conclusions were derived using data from a pig breeding program, we believe that they can be generalized to other animal breeding programs, because preselection presumably has the same effect in any animal breeding program.

## Declarations

### Ethical approval

The data used for this study were collected as part of routine data recording in a commercial breeding program. Samples collected for DNA extraction were used for routine diagnostic purposes of the breeding program. Data recording and sample collection were conducted in line with local laws on protection of animals.

### Availability of data

The data used in the present study were provided by Topigs Norsvin, and are not publicly accessible.

### Authors’ contributions

All authors participated in the conception and the design of the study and of the analysis of the dataset. RB provided the dataset, IJ analysed the dataset and wrote the first draft of the manuscript, and participated in revising the manuscript. All authors read and approved the final manuscript.

## Acknowledgements

The authors thank Marco Bink and Katrijn Peeters from Hendrix Genetics, John Henshall from Cobb Europe, and Chris Schrooten and Gerben de Jong from CRV, for their inputs towards the design of this study.

