## Supplementary material for "Impact of genomic preselection on subsequent genetic evaluations with ssGBLUP - using real data from pigs": Table S3

**Table S3** Estimated additive genetic and residual variances (standard errors in bracket) in the dam line

| Variance/Preselection scenario | With records on animals  in the validation generation | | | Without records on animals  in the validation generation | |
| --- | --- | --- | --- | --- | --- |
|  | Reference^a^ | VGP^b^ | MGP^c^ | Reference/VGP | MGP^c^ |
| *Average daily gain during performance testing* | | | | | |
| Additive genetic | 2542 (72) | 2575 (83) | 2605 (156) | 2374 (80) | 2392 (163) |
| Residual | 5659 (44) | 5343 (50) | 3968 (102) | 5430 (50) | 3860 (109) |
| *Average daily gain throughout life* | | | | | |
| Additive genetic | 844 (24) | 885 (28) | 928 (53) | 835 (27) | 933 (59) |
| Residual | 1871 (14) | 1796 (17) | 1246 (34) | 1825 (17) | 1211 (37) |
| *Backfat thickness* | | | | | |
| Additive genetic | 1.63 (0.03) | 1.68 (0.04) | 1.62 (0.08) | 1.70 (0.04) | 1.73 (0.10) |
| Residual | 1.57 (0.02) | 1.63 (0.02) | 1.55 (0.05) | 1.63 (0.02) | 1.56 (0.06) |
| *Loin depth* | | | | | |
| Additive genetic | 5.90 (0.14) | 5.84 (0.16) | 6.24 (0.33) | 5.78 (0.16) | 6.02 (0.34) |
| Residual | 5.96 (0.07) | 5.93 (0.08) | 5.20 (0.19) | 5.96 (0.08) | 5.25 (0.20) |

^a^ In the reference scenario, the subsequent ssGBLUP evaluation utilized the entire available data until the validation generation

^b^ Validation generation preselection (VGP) scenario, in which all animals in the validation generation without progeny in the data were discarded

^c^ Multi-generation preselection (MGP) scenario, in which all animals in the validation and training generations without progeny in the data were discarded
