## Supplementary material for "Impact of genomic preselection on subsequent genetic evaluations with ssGBLUP - using real data from pigs": Table S2

**Table S2** Estimated additive genetic and residual variances (standard errors in bracket) in the sire line

| Variance/Preselection scenario | With records on animals in the validation generation \| With records on animals in the validation generation | | | Without records on animals in the validation generation \| With records on animals in the validation generation | |
| --- | --- | --- | --- | --- | --- |
|  | Reference^a^ | VGP^b^ | MGP^c^ | Reference/VGP | MGP^c^ |
| *Average daily gain during performance testing* | | | | | |
| Additive genetic | 2369 (101) | 2486 (120) | 2139 (189) | 2357 (118) | 2333 (228) |
| Residual | 7548 (67) | 7526 (80) | 4293 (135) | 7668 (80) | 4254 (160) |
| *Average daily gain throughout life* | | | | | |
| Additive genetic | 1048 (43) | 1102 (51) | 929 (82) | 1057 (50) | 961 (95) |
| Residual | 2939 (27) | 2878 (32) | 1821 (58) | 2922 (32) | 1771 (66) |
| *Backfat thickness* | | | | | |
| Additive genetic | 1.41 (0.04) | 1.46 (0.05) | 1.29 (0.08) | 1.47 (0.05) | 1.38 (0.10) |
| Residual | 1.03 (0.02) | 1.06 (0.02) | 0.95 (0.04) | 1.07 (0.02) | 0.94 (0.05) |
| *Loin depth* | | | | | |
| Additive genetic | 7.96 (0.22) | 7.90 (0.25) | 7.45 (0.49) | 7.96 (0.26) | 7.80 (0.58) |
| Residual | 6.44 (0.11) | 6.40 (0.13) | 6.06 (0.27) | 6.39 (0.13) | 5.90 (0.32) |

^a^ In the reference scenario, the subsequent ssGBLUP evaluation utilized the entire available data until the validation generation

^b^ Validation generation preselection (VGP) scenario, in which all animals in the validation generation without progeny in the data were discarded

^c^ Multi-generation preselection (MGP) scenario, in which all animals in the validation and training generations without progeny in the data were discarded
