## Supplementary material for "Impact of genomic preselection on subsequent genetic evaluations with ssGBLUP - using real data from pigs": Table S1

**Table S1** Estimated heritabilities (diagonal), genetic correlations (below diagonal) and phenotic correlations (above diagonals) for the traits utilized in this study, using the full data from the sire line. Standard errors are in brackets.

| Traits | ADGT | ADGL | Backfat thickness | Loin depth |
| --- | --- | --- | --- | --- |
| ADGT | **0.24 (0.01)** | 0.91 (0.00) | 0.20 (0.01) | -0.17 (0.01) |
| ADGL | 0.92 (0.00) | **0.26 (0.01)** | 0.20 (0.01) | -0.17 (0.01) |
| Backfat thickness | 0.27 (0.02) | 0.32 (0.02) | **0.58 (0.01)** | -0.04 (0.01) |
| Loin depth | -0.29 (0.02) | -0.30 (0.02) | -0.11 (0.02) | **0.55 (0.01)** |

ADGT: Average daily gain during performance testing

ADGL: Average daily gain throughout life
